# Cost Effective Acoustic Monitoring of Biodiversity and Bird Populations in Kenya

**DOI:** 10.1101/072546

**Authors:** Ciira wa Maina, David Muchiri, Peter Njoroge

## Abstract

With the increasing need to effectively monitor a growing number of ecosystems of interest due to risks posed to these ecosystems by human activity and climate change, novel approaches to biodiversity monitoring are needed. In this work we demonstrate the application of low cost acoustic recorders based on the Raspberry Pi microprocessor to biodiversity monitoring. The recorders are capable of capturing audio recordings from which we can compute acoustic indices of biodiversity and identify bird species of interest. We compare the acoustic indices of biodiversity and results of point counts aimed at determining bird species presence and find that the acoustic complexity index has a significant positive correlation to point count results. In addition, we show that the presence of the Hartlaub’s Turaco, a ubiquitous species in montane forests in Kenya with a distinct call, can be automatically determined using recordings obtained using our setup. Montane species are of interest for long-term automatic monitoring since they are particularly vulnerable to the effects of climate change. Our system is able to deal with the large amounts of data generated by the acoustic recorders. The automatic screening of approximately five hours of recordings for presence of the Hartlaub’s Turaco call is achieved in approximately three minutes representing a large time saving that makes use of audio recordings for species identification feasible.

## Introduction

With increasing demands on natural resources, there is a growing need to ensure that the exploitation of these resources is environmentally sustainable. In addition, several ecosystems are under threat from human encroachment and climate change. There is therefore a need to monitor the biodiversity of various ecosystems to ensure that corrective measures are initiated if species loss or habitat destruction is detected. Traditional methods of biodiversity assessment involve survey of plant and animal species in habitats of interest by trained experts [1]. These methods are costly and time consuming and cannot be applied on the scale necessary to monitor all ecosystems of interest. Techniques such as rapid biodiversity assessment (RBA) aim to reduce the amount of effort required to survey an area by focusing on a few indicator species that serve to gauge the health of a particular ecosystem [2]. RBA still requires trained experts who can identify members of the indicator species and is therefore difficult to scale.

The challenges in carrying out traditional biodiversity assessment have led to interest in bioacoustic approaches to biodiversity assessment [3,4]. Using acoustic recordings obtained from an ecosystem of interest, methods to infer the species richness and diversity of the ecosystem from which the recordings have been obtained have been developed. A number of acoustic indices such as acoustic entropy (AE) and the acoustic complexity index (ACI) can be readily computed from audio recordings and have been shown to be correlated to species richness [3,5]. These acoustic indices do not rely on any information about the identity of vocalizing species. However, using these recordings, systems to identify vocalizing species can be developed and these have been shown to be effective in surveying species of interest.

The use of acoustic recordings in biodiversity assessment has a number of advantages over traditional survey techniques. Firstly, the audio recordings can be archived to serve as a permanent record of the state of the ecosystems at the time of the survey. Secondly, experts are not needed to collect the data as the recordings can be collected by people trained to handle the recording equipment only. Thirdly, the acoustic recordings can be used to perform analysis at various levels starting from a large scale biodiversity assessment to a survey of a particular species [6]. Fourth, the recording equipment is not intrusive. Fifth, acoustic recordings can be used for long-term monitoring of ecosystems of interest. This has the potential to track individual species over long periods and to determine species specific behavior such as migration patterns. For example in [7], Frommolt *et al.* demonstrate the use of acoustic recordings obtained over five years to track the population and distribution of Eurasian bittern (*Botaurus stellaris*). Similar long-term efforts have been used to monitor sperm whales using acoustic recordings [8].

Despite these advantages, a number of shortcomings remain. Firstly, recording equipment can be costly to obtain and setup. Second, audio recording generates a large amount of data which can be time consuming to analyze. Third, it is not straight forward to determine the number of vocalizing individuals from an acoustic recording. Fourth, the reliance of the method on acoustic signals may lead to an overestimation of the abundance of vocalizing species while the abundance of species that do not vocalize or vocalize rarely will be underestimated. As a result, efforts to make bioacoustic approaches more accurate, cost effective and less time consuming are important and deserve the attention of the research community. In [9], the authors describe the use of a low cost audio recorder to monitor diverse ecosystems. In [10], the authors use low cost audio recorders to assess agroforestry systems. In addition to testing low cost devices for use in biodiversity assessment. Systems to analyze the large datasets generated by acoustic recorders are necessary. In [11] the authors describe a system to screen large datasets for the vocalization of a particular species, the Screaming Piha (*Lipaugus vociferans*) in a tropical forest in French Guiana.

In this work we develop and test a low cost acoustic monitoring system based on the Raspberry Pi microprocessor for use in acoustic biodiversity monitoring. The Raspberry Pi is a low cost programmable microprocessor with much of the functionality of a modern computer and this makes the devices suitable for both recording and processing of the recordings. We assess the utility of the recordings obtained by comparing acoustic indices of biodiversity computed from these recordings to the results of bird point count data obtained from the same area. In addition, we use the recordings to detect the presence of the Hartlaub’s Turaco (*Tauraco hartlaubi*), a ubiquitous species in montane forests in Kenya with a distinct call.

Montane species in tropical ecosystems are some of the most vulnerable to the effects of climate change [12]. It is predicted that these birds will be forced to move to higher altitudes as temperatures rise and this could lead to extinction of several species. It is therefore important to initiate long-term monitoring efforts for these birds so that any reduction in their range is detected. In Kenya, the Hartlaub’s Turaco is of special conservation interest because it is predicted that it will lose more than 50% of its range due to climate change [13]. The work presented here provides technology that could aid long-term monitoring efforts of this species which could in turn provide a measure of the effects of climate change.

## Materials and Methods

### Ethics Statement

Data for this study were collected in a non-invasive manner and therefore no ethical approval was necessary.

### Study Area

The study was conducted at the Dedan Kimathi University Wildlife Conservancy (DeKUWC) located at 0°23’17.0”S 36°57’43.2”E at an elevation of approximately 1800m. The conservancy covers an area of 120 Acres with three ecological zones namely open grassland, undisturbed indigenous forest and aquatic zones. The DeKUWC is located in the central part of Kenya and receives approximately 1000mm of rainfall annually. This is in two rainy seasons of mid March to May and October to November. The conservancy is part of the Mount Kenya ecosystem with the Kabiruini forest bordering it to the North. The Kabiruini forest has suffered from human encroachment with quarrying and cattle grazing occurring within the forest. To the South, the conservancy is bordered by human settlements and a major highway (see the map in Figure 3).

### Equipment

The audio recordings were collected using a cheap microphone connected to a Raspberry Pi (RPi). The RPi is a cheap credit card sized microprocessor with functionality similar to an ordinary computer. The RPi runs the Raspbian operating system - Debian Wheezy which is quite similar to Ubuntu. To make the recordings, we use the open source sound processing software SoX. Installation instructions are available on the project website http://sox.sourceforge.net/. The Raspberry Pi was programmed to obtain one minute recordings every five minutes.

It was found that this setup was capable of recording audio signals that were of good enough quality to recognize the species vocalizing in the recording. Also, it was possible to use these simple microphones to detect noise sources such as gunshots and cow bells. These sounds are of interest in ecological monitoring because they are indicative of human encroachment into wildlife habitats. Figure 1 shows the acoustic sensor system in the lab and deployed in the field. Figure 2 shows the spectrogram of a typical recording. There are a number of bird species vocalizing in the recording (labeled A-D) and gunshot noise from the neighbouring Kiganjo Police Training college (E).

**Figure 1.**
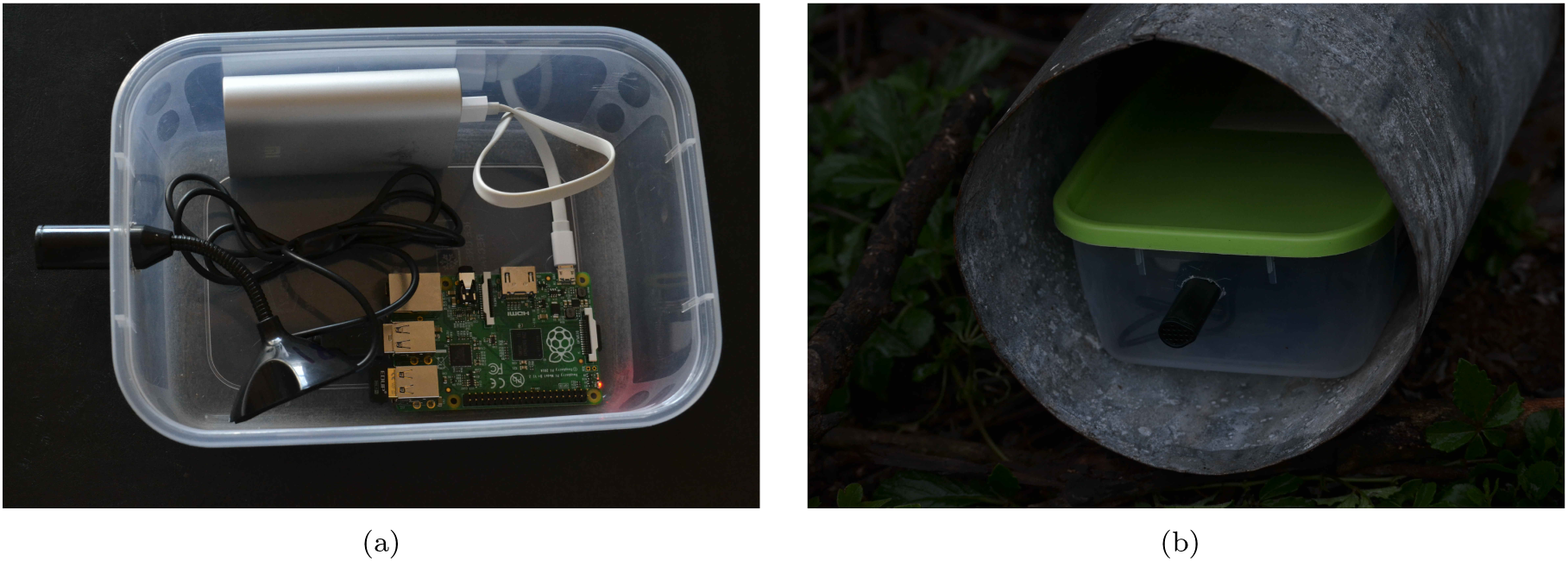
The Raspberry Pi based acoustic sensor system (a) and the system deployed at the DeKUT conservancy (b).

**Figure 2.**
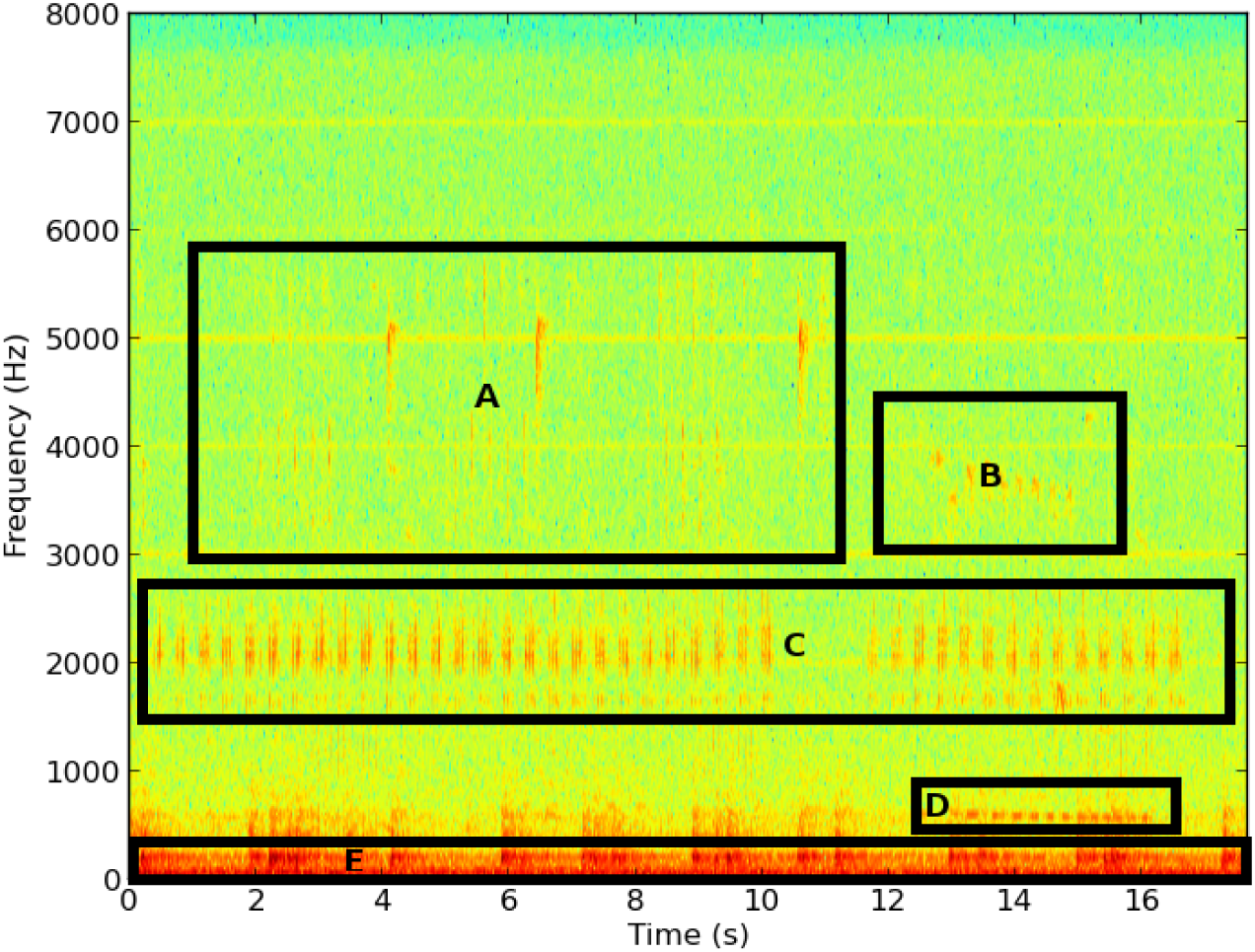
The spectrogram of a typical recording obtained at the DeKUT conservancy. There are a number of bird species vocalizing in the recording (labeled A-D) and gunshot noise from the neighbouring Kiganjo Police Training college (E).

### Acoustic Survey Protocol

We performed point counts at twenty locations within the Dedan Kimathi University of Technology wildlife conservancy (DeKUWC) on two different days: 5th January, 2016 (10am to 12noon) and 28th January, 2016 (8am to 10am) (ten on each day). The points were separated by approximately 40 meters and birds seen or heard and judged to be within 20m of each location were recorded for ten minutes. The 20 point count locations are labeled A-T. We used four acoustic recorders and these were left at locations near some of the point count locations. A total of eight locations were sampled, four on each day. The acoustic recorder locations are labeled 1-8. The locations are shown on the map in Figure 3. An interactive version of this map is available via this link: 

~~~
https://goo.gl/BRn95u
~~~

. The recorders were left at these points for approximately 28 hours and were programmed to record for one minute at five minute intervals. This produced approximately 340 minute long recordings per site. We set the sampling rate of the recorders to 16kHz at 16 bit resolution.

**Figure 3.**
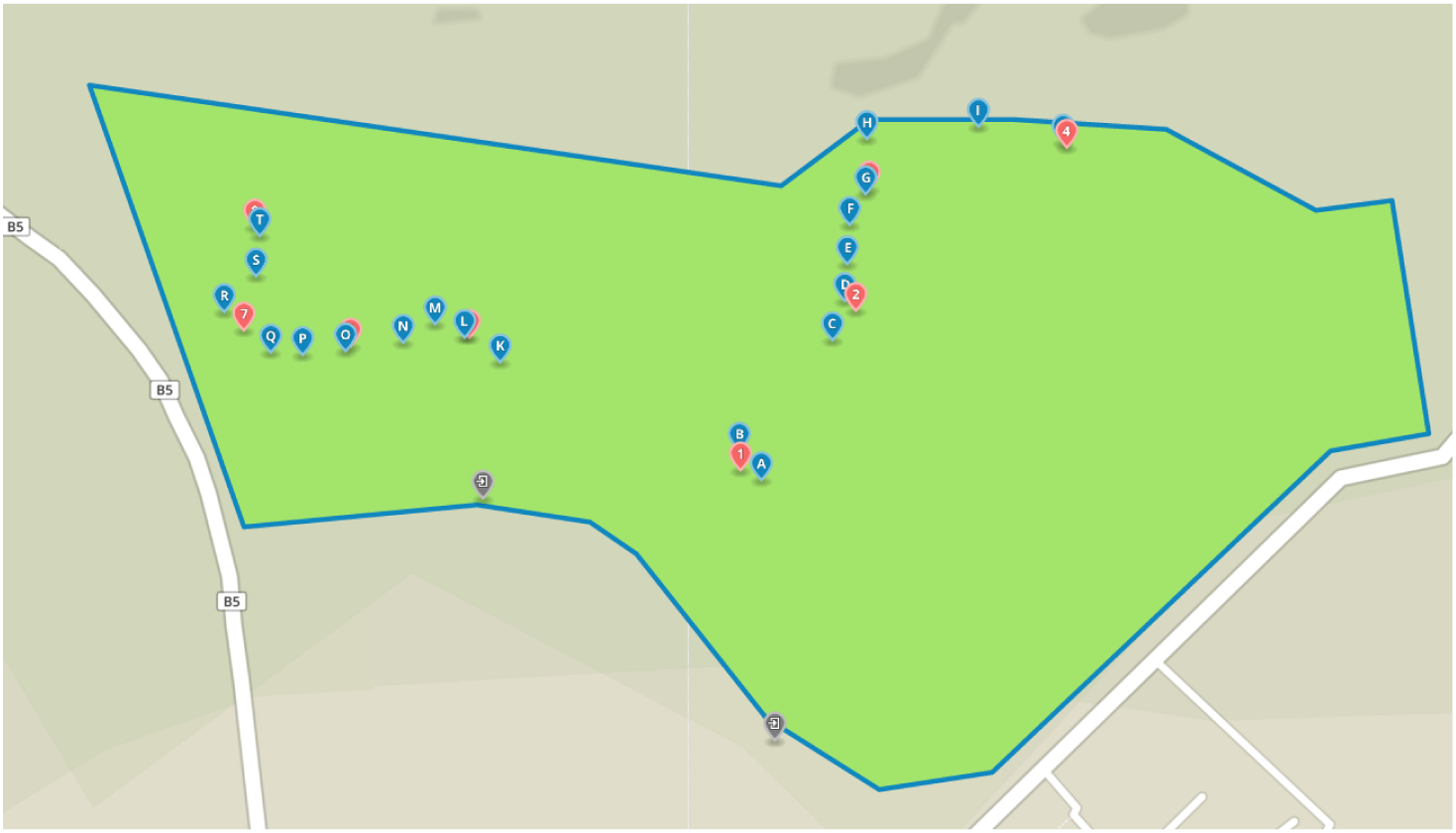
A map of the DeKUWC with locations of the point counts and acoustic recorders indicated. The 20 point count locations are labeled A-T and the acoustic recorder locations are labeled 1-8.

### Signal Processing

In this work we aim to infer both the species richness of a location and the presence of a particular bird species from audio recordings. This requires a number of signal processing operations to be performed on the recordings.

#### Acoustic Indices of Biodiversity

Acoustic indices of biodiversity aim to infer the species richness of an ecosystem using audio recordings obtained from the ecosystem of interest. In this work we focus on two indices namely acoustic entropy [3] and the acoustic complexity index [5].

- **Acoustic Entropy**: Entropy is a measure of the uncertainty associated with a random variable [14]. It can be viewed as a measure of information obtained when the value of the random variable is observed. In acoustic recordings the notion of entropy is related to the acoustic activity in a recording. If there is little acoustic activity in a recording, for example in a quiet room, this corresponds to a low acoustic entropy. If there is significant and varied acoustic activity, for example in a noisy room, this corresponds to a high acoustic entropy. It has been shown that the acoustic entropy of a recording is correlated to species richness and the number of species vocalizing in the recording [3, 4, 15].
- **Acoustic Complexity Index**: The acoustic complexity index (ACI) is based on the observation that spectrograms of audio signals from natural environments exhibit sudden changes in intensity of signal energy at particular frequency bins. This index quantifies the changes in this energy from one frame to the next.

#### Acoustic Features

In order to identify a particular bird species from its vocalization, in this case the Hartlaub’s Turaco (*Tauraco hartlaubi*), we need to extract features from the recording which help distinguish the bird. These features are then used to train models that will be used in our classifiers. Figure 4(a) shows a typical spectrogram obtained from a recording containing the Hartlaub’s Turaco vocalization while Figure 4(b), shows a typical recording without the Hartlaub’s Turaco call. The spectral signature of the Hartlaub’s Turaco call is clearly visible.

**Figure 4.**
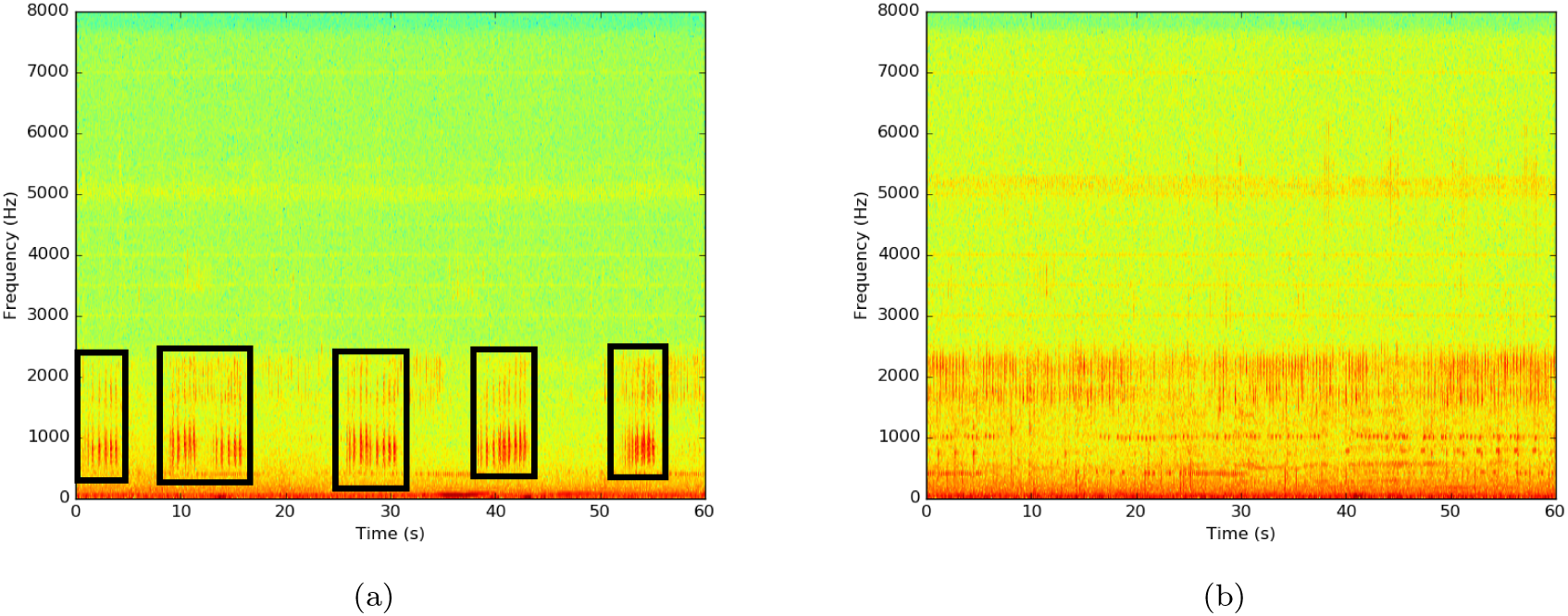
A typical spectrogram obtained from a recording containing the Hartlaub’s Turaco vocalization. Regions with the vocalization are indicated using the black rectangles (a). A typical spectrogram obtained from a recording without the Hartlaub’s Turaco vocalization (b).

Mel frequency cepstral coefficients (MFCCs) are commonly used in speech processing applications and a number of authors have used them to recognize bird species from recordings [16]. In this work we also use simpler spectral features which have been used in music genre [17,18] and acoustic scene classification [19,20] and compare their performance to MFCCs. These features are:

1. Band Energy Ratio: This is the ratio of the total energy in a particular spectral band to the total energy of a frame. We use six logarithmically spaced spectral bands.
2. Spectral Flux: This is the squared difference between the normalized magnitude spectra of successive frames.
3. Spectral centroid and bandwidth: The spectral centroid is a measure of the frequency around which spectral energy is centered while bandwidth measures the dispersion of spectral energy around this centroid frequency.
4. Spectral Rolloff: This is the the frequency below which a given percentage of the spectral energy is contained. In this work we use 85%.

These features are computed from the spectrogram of the acoustic recording. To obtain the spectrograms, we divide the signal into frames of 512 samples each (32 ms at 16kHz) with 50% overlap and compute the magnitude of the short time Fourier transform (STFT) of each frame. The STFT is computed using the fast Fourier transform (FFT).

#### Classification

In order to detect the presence of the Hartlaub’s Turaco, we train two Gaussian mixture models (GMMs) using the spectral features and MFCCs obtained from the acoustic recordings. One model is trained using positive examples of recordings containing the Hartlaub’s Turaco call and the other is trained using recordings which do not contain the Hartlaub’s Turaco call.

Given a set of features χ = {x_1_, …, x_*N*_} obtained from a recording, we would like to test which hypothesis is true:

- *H*_0_ : The recording does not contain the Hartlaub’s Turaco call
- *H*_1_ : The recording contains the Hartlaub’s Turaco call

Mathematically we would like to determine which of the following is true:

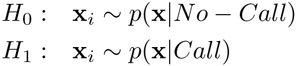

Where *p*(x|.) is the probability distribution of the features. The distributions are assumed to be GMMs and take the following mathematical form

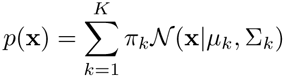

where 𝓝(x|μ_*k*_, Σ_*k*_) is a Gaussian distribution with mean μ_*k*_ and covariance Σ_*k*_ and the mixture coefficients π_*k*_ satisfy the condition 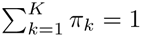.

The parameters of these distributions {π_*k*_, μ_*k*_, Σ_*k*_} are learnt from the training data using the expectation-maximization algorithm. The GMMs are initialized by applying K-means clustering to the training data with the number of mixture coefficients determined by cross-validation. The models and classifiers were implemented using Bob, an open source machine learning toolkit written in python [21].

To decide between the two hypotheses, we compute the log likelihood ratio (LLR) of the two models and compare the ratio to a threshold λ. That is we compute

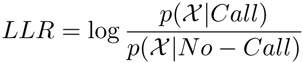

and compare this quantity to λ. If *LLR* > λ we assume that *H*_1_ is true and if *LLR* < λ we assume that *H*_0_ is true.

#### Counting Hartlaub’s Turaco Call Events

In addition to detection of recordings containing the call of the Hartlaub’s Turaco, we are interested in counting the number of calls in a particular recording. The call rate can be used to infer the number of individuals present near a recorder. If the baseline call rate in a region is known, increase in this call rate could indicate the presence of additional individuals in the area. Also, changes in the call rate could indicate behavioral changes due to breeding or seasonal change. Here, the call rate will be used to indicate presence of the Hartlaub’s Turaco near a recorder and also determine the calling behavior. That is, we will be able to determine when the Hartlaub’s Turaco calls actively.

In this work we use an approach similar to that used in note onset detection in music [22]. Here we first derive a detection function from the acoustic recording. Peaks in the detection function correspond to locations of the Hartlaub’s Turaco call. The detection function we use is derived from the recordings using the following steps.

1. Divide the signal into frames of 512 samples each (32 ms at 16kHz) with 50% overlap and extract the features used to classify the Harlaub’s Turaco call.
2. Compute the log likelihood ratio of each frame using the models trained to identify turaco calls and no turaco calls.
3. Smooth this log likelihood ratio by computing the average of log likelihood of the 300 previous blocks. This is so that we only consider blocks in a 5 second window which is the approximate duration of a call. The smoothed LLR will serve as the detection function.
4. Compare the smoothed LLR to a threshold and determine the regions where the smoothed LLR is above the threshold. These regions correspond to the detected calls.

### Dataset

The dataset consists of approximately 2700 minute long recordings obtained from the four RPi recorders on the two study days: 5th and 28th January, 2016. This dataset was used to

1. Compare the results from point counts and audio recorders in determining bird species presence.
2. Compare the use of acoustic indices of biodiversity and results from point counts in determining species richness.
3. Develop and test an automatic species recognizer for the Hartlaub’s Turaco call for use in screening large datasets.

#### Hartlaub’s Turaco Dataset

From the recordings obtained from recorder location 4 on 5th January, 2016, 12 recordings were chosen which contained vocalizations of the Hartlaub’s Turaco and 21 recordings which did not contain the call. Each of the recordings was split into six, ten second long recordings for use in training and testing our single species classifier. Thus in total there were 198 recordings. Of these 60 had Hartlaub’s Turaco calls since some segments obtained from recordings containing the calls did not have calls.

## Results

### Comparison of Point Counts and Acoustic Recordings

We sought to compare the number of species identified via point counts and identification of vocalizations by listening to the recordings. Table 1 shows the result of the point count on 5th January, 2016 and the species identified from recording their vocalisations. We see that of the 40 species identified, 26 were identified during the point count, 12 were identified during both the point count and using the recordings while 14 were identified using the recordings only.

**Table 1.** Results of the point count on 5th January, 2016 and the species identified from recordings obtained at four locations near point count locations.

| # | Species | Point Count Location |  |  |  |  |  |  |  |  |  | NOI | Acoustic Recorder Location (Nearest Point Count Location) |  |  |  |
| --- | --- | --- | --- | --- | --- | --- | --- | --- | --- | --- | --- | --- | --- | --- | --- | --- |
|  |  | A | B | C | D | E | F | G | H | I | J |  | 1(B) | 2(D) | 3(G) | 4(J) |
| 1 | Red-headed Weaver | 2 |  |  |  |  |  |  |  |  |  | 2 |  |  |  |  |
| 2 | African Paradise Flycatcher | 1 | 1 |  |  |  |  |  |  |  |  | 2 |  | ✓ |  | ✓ |
| 3 | Cinnamon-chested Bee-eater | 1 |  |  |  |  |  |  |  |  |  | 1 |  |  |  |  |
| 4 | Common Bulbul |  | 3 | 1 | 1 |  | 1 |  | 1 | 1 | 3 | 11 | ✓ | ✓ | ✓ | ✓ |
| 5 | Black-collared Apalis |  | 1 |  |  |  |  |  |  |  |  | 1 |  |  |  |  |
| 6 | Black-backed Puffback |  | 1 |  |  |  |  |  |  |  |  | 1 | ✓ | ✓ |  | ✓ |
| 7 | Yellow-whiskered Greenbul |  |  | 1 | 1 |  |  | 1 | 1 |  | 1 | 5 | ✓ | ✓ | ✓ | ✓ |
| 8 | Variable Sunbird |  |  | 1 | 1 | 1 |  |  |  |  | 1 | 4 |  | ✓ |  |  |
| 9 | Tropical Boubou |  |  | 1 |  |  |  |  |  |  |  | 1 | ✓ | ✓ | ✓ | ✓ |
| 10 | African Golden-breasted Bunting |  |  | 1 |  |  |  |  |  |  |  | 1 |  |  |  |  |
| 11 | Grey-backed Camaroptera |  |  | 1 |  |  | 1 | 1 |  |  |  | 3 | ✓ | ✓ | ✓ | ✓ |
| 12 | Eurasian Blackcap |  |  |  | 1 |  |  |  |  |  |  | 1 |  |  |  |  |
| 13 | Yellow-breasted Apalis |  |  |  |  | 1 |  |  |  |  |  | 1 |  | ✓ |  |  |
| 14 | Ring-necked Dove |  |  |  |  |  | 1 |  |  |  |  | 1 |  |  |  |  |
| 15 | Yellow-rumped Tinkerbird |  |  |  |  |  |  | 2 |  | 1 |  | 3 | ✓ | ✓ | ✓ | ✓ |
| 16 | Augur Buzzard |  |  |  |  |  |  |  | 1 |  |  | 1 |  |  |  |  |
| 17 | African Dusky Flycatcher |  |  |  |  |  |  |  | 1 |  |  | 1 |  |  |  |  |
| 18 | Grey Apalis |  |  |  |  |  |  |  |  | 1 |  | 1 |  |  |  |  |
| 19 | Spectacled Weaver |  |  |  |  |  |  |  |  | 1 |  | 1 |  |  |  |  |
| 20 | Tambourine Dove |  |  |  |  |  |  |  |  |  | 1 | 1 | ✓ | ✓ | ✓ | ✓ |
| 21 | Eastern Double-collared Sunbird |  |  |  |  |  |  |  |  |  | 1 | 1 |  |  |  |  |
| 22 | Chinspot Batis |  |  |  |  |  |  |  |  |  | 1 | 1 |  | ✓ |  | ✓ |
| 23 | Southern Black Flycatcher |  |  |  |  |  |  |  |  |  | 1 | 1 |  |  |  |  |
| 24 | Speckled Mousebird |  |  |  |  |  |  |  |  |  | 4 | 4 |  |  |  |  |
| 25 | Olive Thrush |  |  |  |  |  |  |  |  |  | 1 | 1 | ✓ |  |  |  |
| 26 | African Grey Flycatcher |  |  |  |  |  |  |  |  |  | 1 | 1 |  |  |  |  |
| 27 | Hadada Ibis |  |  |  |  |  |  |  |  |  |  | - | ✓ |  |  |  |
| 28 | Hartlaub's Turaco |  |  |  |  |  |  |  |  |  |  | - | ✓ | ✓ | ✓ | ✓ |
| 29 | Pied Crow |  |  |  |  |  |  |  |  |  |  | - | ✓ |  |  |  |
| 30 | Abyssinian Crimsonwing |  |  |  |  |  |  |  |  |  |  | - | ✓ |  |  |  |
| 31 | Black-headed Oriole |  |  |  |  |  |  |  |  |  |  | - | ✓ |  |  |  |
| 32 | Silvery-cheeked Hornbill |  |  |  |  |  |  |  |  |  |  | - | ✓ |  |  |  |
| 33 | Brown Woodland Warbler |  |  |  |  |  |  |  |  |  |  | - | ✓ | ✓ | ✓ |  |
| 34 | Fork-tailed Drongo |  |  |  |  |  |  |  |  |  |  | - | ✓ | ✓ |  |  |
| 35 | Spot-flanked Barbet |  |  |  |  |  |  |  |  |  |  | - |  | ✓ |  |  |
| 36 | Ruppel's Robin-Chat |  |  |  |  |  |  |  |  |  |  | - |  | ✓ |  |  |
| 37 | Collared Sunbird |  |  |  |  |  |  |  |  |  |  | - |  |  | ✓ | ✓ |
| 38 | Black Cuckoo |  |  |  |  |  |  |  |  |  |  | - |  |  | ✓ |  |
| 39 | Olive Sunbird |  |  |  |  |  |  |  |  |  |  | - |  |  |  | ✓ |
| 40 | Black-throated Wattle-eye |  |  |  |  |  |  |  |  |  |  | - |  |  |  | ✓ |
|  | **Number of Species** | 3 | 4 | 6 | 4 | 2 | 3 | 3 | 4 | 4 | 10 | - | 16 | 16 | 10 | 13 |

Table 2 shows the result of the point count on 28th January, 2016 and the species identified from recording their vocalizations. We see that of the 35 species identified, 17 were identified during the point count, 9 were identified during both the point count and using the recordings while 18 were identified using the recordings only.

**Table 2.**
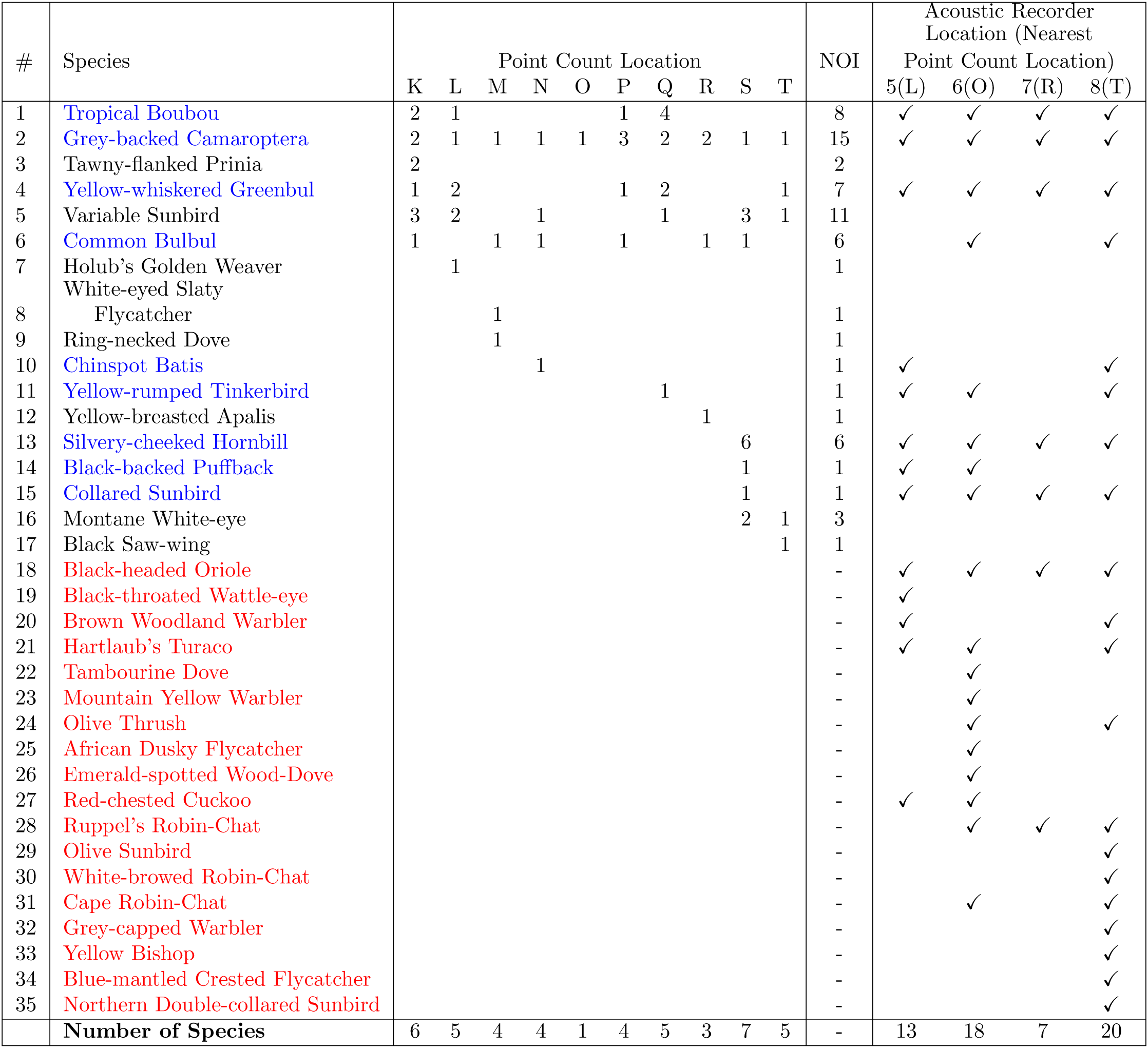
Results of the point count on 28th January, 2016 and the species identified from recordings obtained at four locations near point count locations.

A total of 54 bird species were observed during the study. A list of these bird species is contained in the supplementary material. We include the common name, scientific name and a four letter code used to identify the birds.

Species identified by both point counts and acoustic recordings are highlighted in blue while species identified by acoustic recordings only are highlighted in red. NOI is the Number of Individuals recorded.

### Acoustic Indices of Biodiversity

Using the recordings obtained at each of the recording locations, we computed the acoustic entropy (AE) and acoustic complexity index (ACI) and plotted it as a function of time. The acoustic indices of biodiversity were computed using the R package seewave [23]. Figure 5(a) shows a plot of the AE as a function of time at the third location (R3) from 12:00 on January 5, 2016 to 15:15 on January 6, 2016. Figure 5(b) shows a plot of the ACI at the same location over the same time. We see that the ACI is significantly lower at night with a spike at dawn clearly visible.

**Figure 5.**
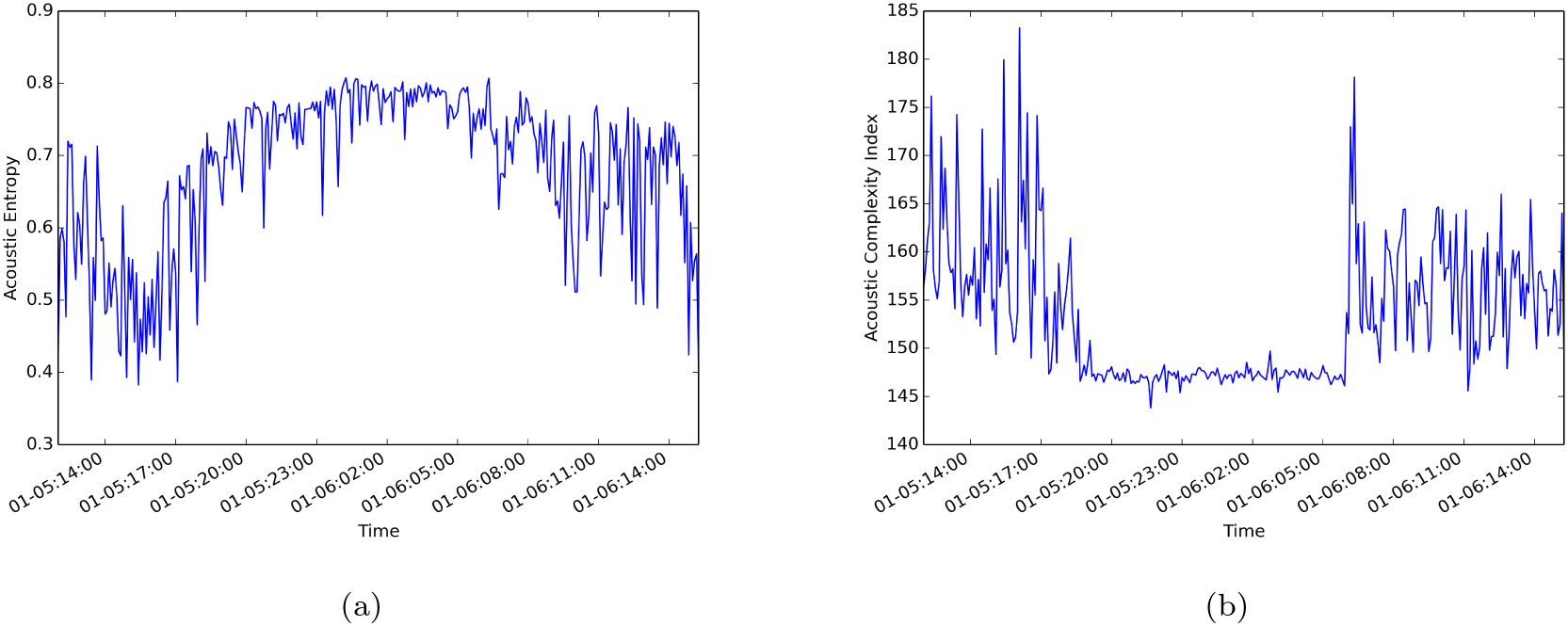
The AE as a function of time at the third location from 12:00 on January 5, 2016 to 15:15 on January 6, 2016 (a) and a plot of the ACI at the same location over the same time (b).

### Comparison of Acoustic Indices of Biodiversity and Point Counts

To investigate the relationship between the acoustic indices of biodiversity namely the acoustic entropy (AE) and acoustic complexity index (ACI) and point counts, we computed the mean ACI and AE at the eight recorder locations 1-8 during the day (05:30-18:30) and compared it to the number of species identified using the acoustic recordings and those observed at nearby point count locations. Table 3 shows the ACI, AE, number of bird species identified using the acoustic recordings (#AR) and the number of bird species observed at nearby point count locations (#PC) for each of the recorder locations 1-8. Table 4 shows the Spearman’s correlation and corresponding p-value between each of these four quantities. From these results we see that a significant positive correlation of 0.73 exists between the ACI and the number of bird species identified during the point count.

**Table 3.** The ACI, AE, number of bird species identified using the acoustic recordings and the number of bird species observed at nearby point count locations for each of the recorder locations 1-8.

| Acoustic recorder location<br>(Nearby point count location) | ACI | AE | #AR | #PC |
| --- | --- | --- | --- | --- |
| 1 (B) | 153.05 | 0.6245 | 16 | 4 |
| 2 (D) | 162.30 | 0.6839 | 16 | 4 |
| 3 (G) | 157.20 | 0.6304 | 10 | 3 |
| 4 (J) | 165.54 | 0.6919 | 13 | 10 |
| 5 (L) | 162.31 | 0.5937 | 13 | 5 |
| 6 (O) | 154.53 | 0.6361 | 18 | 1 |
| 7 (R) | 150.85 | 0.5784 | 7 | 3 |
| 8 (T) | 157.72 | 0.6360 | 20 | 5 |

**Table 4.** The Spearman’s correlation and corresponding p-value between each of the four variables: ACI, AE, number of bird species identified using the acoustic recordings (#AR) and the number of bird species observed at nearby point count locations (#PC) for each of the recorder locations 1-8.

|  | ACI | AE | #AR | #PC |
| --- | --- | --- | --- | --- |
| ACI | 1 |  |  |  |
| AE | 0.66 (0.076) | 1 |  |  |
| #AR | 0.16 (0.701) | 0.42 (0.30) | 1 |  |
| #PC | **0.73 (0.041)** | 0.51 (0.202) | -0.03 (0.942) | 1 |

### Recognition of The Hartlaub’s Turaco Call

During training and testing, we divide the signal into frames of 512 samples each (32 ms at 16kHz) with 50% overlap and compute the magnitude of the STFT of each frame and extract the features described namely band energy ratio, spectral flux, spectral centroid, bandwidth and spectral rolloff. We used a variation of the spectral flux by computing the flux in each of the six spectral bands. This resulted in a 15-dimensional feature vector. In addition, we computed 19-dimensional MFCCs. To determine the appropriate number of coefficients for both the spectral feature and MFCC models, we used 5-fold cross-validation and computed the receiver operating characteristic (ROC) for models with 2, 4, 8, 16, 32, 64, and 128 mixture coefficients. For each of these models we computed the average area under the ROC curve (AUC). The results are shown in Table 5.

**Table 5.** The average area under the ROC curve for different models.

| Features | Number of Mixtures |  |  |  |  |  |  |
| --- | --- | --- | --- | --- | --- | --- | --- |
|  | 2 | 4 | 8 | 16 | 32 | 64 | 128 |
| Spectral Features | 0.9369 | 0.9622 | 0.9706 | 0.9788 | 0.9771 | **0.9836** | 0.9830 |
| MFCCs | 0.9652 | 0.9878 | 0.9906 | 0.9866 | **0.9921** | 0.9911 | 0.9888 |

Based on these results, we trained a GMM model with 64 mixture coefficients for the the spectral features and a GMM model with 32 mixture coefficients for the MFCCs using all the training data. This resulted in a model with an AUC of 0.995 for the spectral features and 0.998 for the MFCCs. The thresholds for the classifiers were chosen to achieve the true positive rates and false positive rates shown in Table 6.

**Table 6.** Description of the final models for both spectral features and MFCCs.

| Feature | Number of Mixtures | AUC | TPR | FPR |
| --- | --- | --- | --- | --- |
| Spectral Features | 64 | 0.995 | 0.983 | 0.022 |
| MFCCs | 32 | 0.998 | 0.983 | 0.007 |

Code and data to reproduce these experiments is available on 

~~~
Github https://github.com/ciiram/HTuraco
~~~

.

### Screening The Dataset

We run the spectral feature and MFCC based GMM models described in Table 6 on all the recordings obtained at recorder location 4 to determine which files had the Hartlaub’s Turaco call. For each frame, we compute the log likelihood ratio. As our detection function, we use the average log likelihood ratio of the previous 300 frames (approximately 5 seconds) to classify the recording (see Figure 7). If the mean of the detection function is above the threshold, the recording is classified as having the Hartlaub’s Turaco call. Of the 331 files screened, 102 were classified as having putative Hartlaub’s Turaco calls by both the spectral feature and MFCC based GMM models. Each one minute file was processed in approximately 0.5 seconds. In addition, the two models only differed in 25 cases representing an agreement of 92.4%.

Examination of spectrograms obtained from the 102 recordings classified as having the Hartlaub’s Turaco call showed that only 3 recordings were erroneously classified. Also, examination of spectrograms from the 25 files where the spectral feature and MFCC based GMM models differed revealed that the causes of error included:

- The Hartlaub’s Turaco call was too faint.
- Noise sources mistaken for the Hartlaub’s Turaco call.
- Other bird species vocalizations mistaken for the Hartlaub’s Turaco call.

For these files, the result of the MFCC based model was correct in 20 cases while the spectral feature based model was correct in only 5 cases.

Figure 6(a) shows the spectrogram of one of the files classified as containing the call by both the spectral feature and MFCC based GMM models. It is the spectrogram of the recording obtained at 06:15 on 6th January, 2016. Figure 6(b) shows the spectrogram of one of the files classified as not containing the call. It is the spectrogram of the recording obtained at 10:55 on 6th January, 2016.

**Figure 6.**
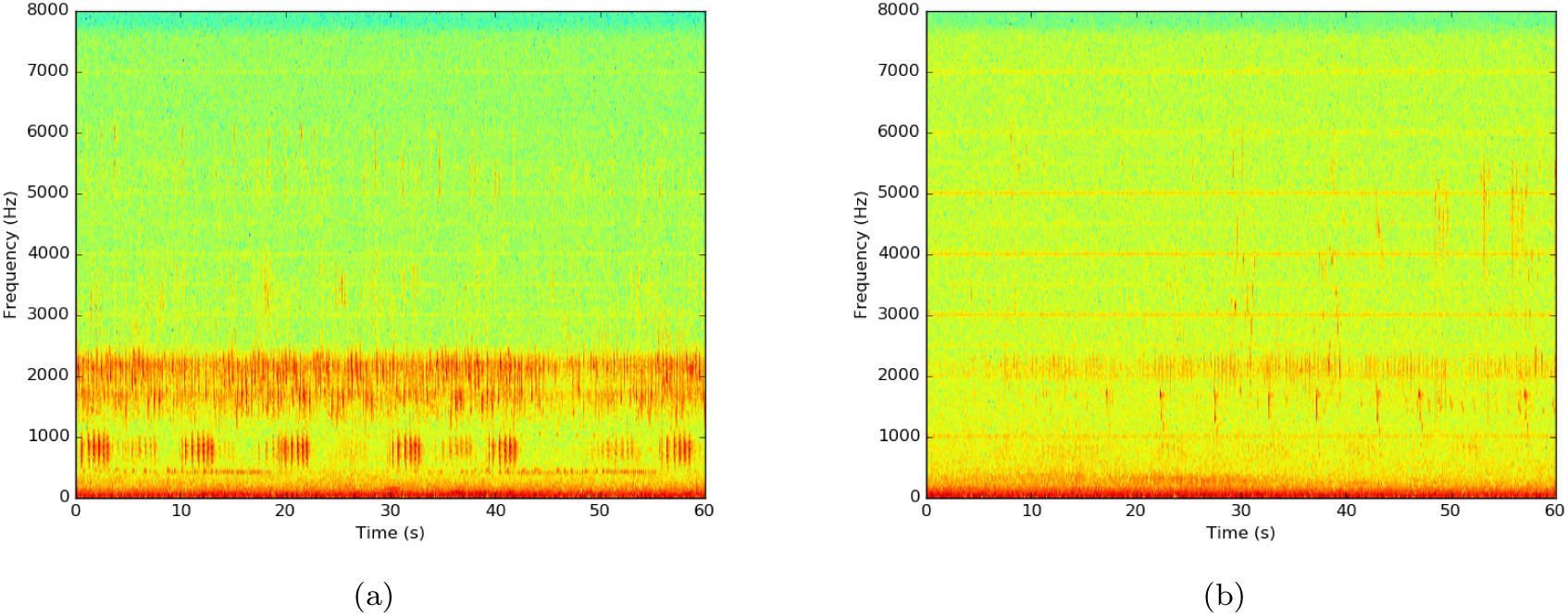
The spectrogram of one of the files classified as containing the Hartlaub’s Turaco call (a). It is the spectrogram of the recording obtained at 06:15 hrs on 6th January, 2016. The spectrogram of one of the files classified as not containing the Hartlaub’s Turaco call (b). It is the spectrogram of the recording obtained at 10:55 hrs on 6th January, 2016.

We also compute the number of Turaco calls detected in each one minute file. To do this, the spectrograms corresponding to the 102 files found to have putative Hartlaub’s Turaco calls by both the spectral feature and MFCC based GMM models were examined and the number of calls in each file noted. Using this as ground truth data, a suitable threshold was chosen to minimize the error in the number of detected calls. For these experiments we used the models trained using MFCCs.

Figure 7 shows the spectrogram of an audio recording containing the Hartlaub’s Turaco call (top panel), the detection function used to extract call locations (middle panel) and the detected calls (bottom panel). From this we see that the detection function peaks at the end of the call and comparing the detection function to a threshold is able to detect the peaks of this function. In this case the four call events are detected.

**Figure 7.**
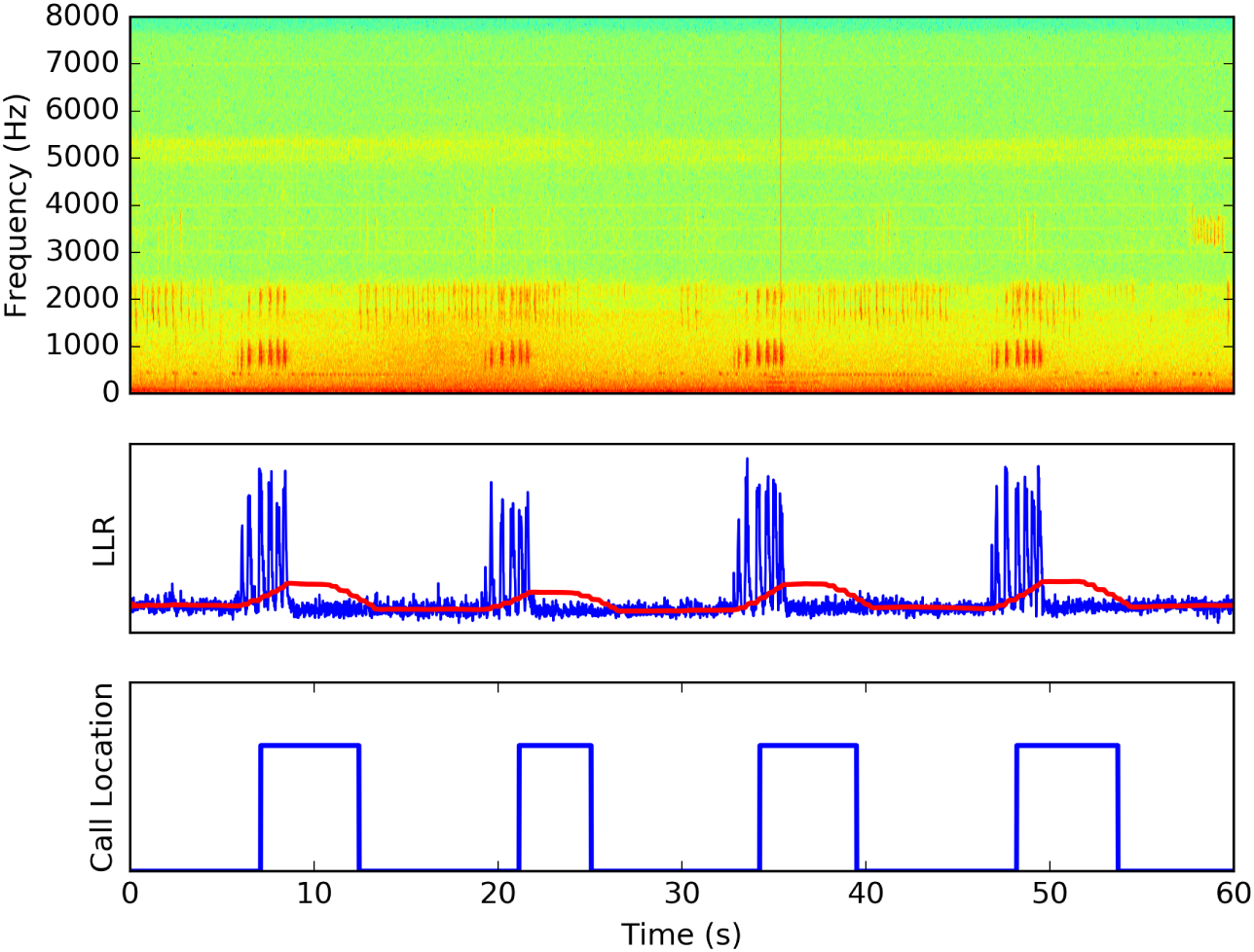
The spectrogram of an audio recording containing the Hartlaub’s Turaco call (top panel), the detection function used to extract call locations (middle panel) and the detected calls (bottom panel). The middle panel shows the log likelihood ratio of each frame (blue curve) and the smoothed log likelihood ratio using a 300 block window (red curve). The smoothed log likelihood ratio is the detection function.

Figure 8 shows a plot of the number of calls per minute detected at recorder location 4 between 12:40 on January 5th, 2016 and 16:10 on January 6th, 2016. We see that calls are detected throughout the day at this location with the latest call detected at 18:45 and the earliest call detected at 06:00.

**Figure 8.**
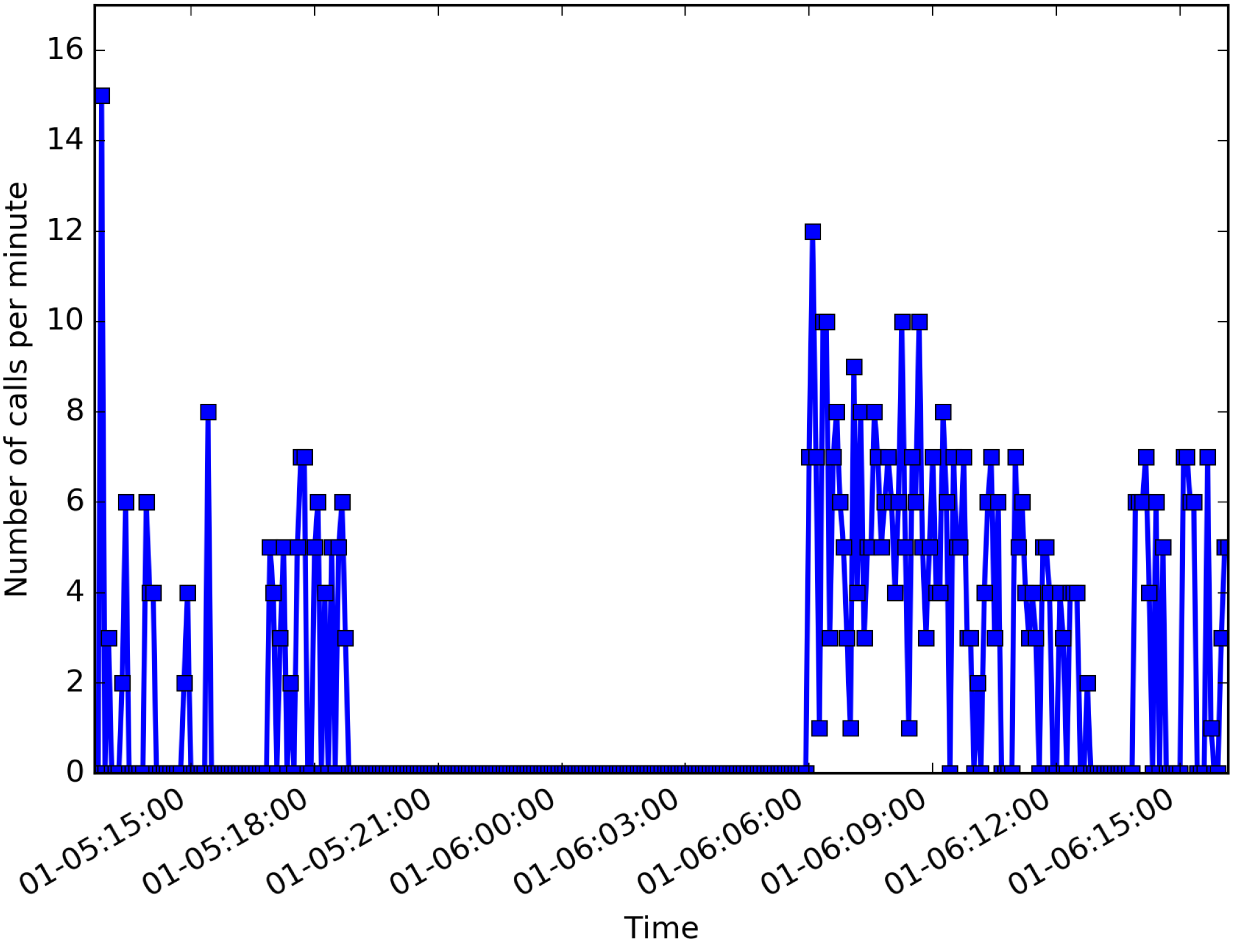
Number of Hartlaub’s Turaco calls per minute detected at recorder location 4 between 12:40 on January 5th, 2016 and 16:10 on January 6th, 2016.

## Discussion

From the results obtained in this work, we see that acoustic indices of biodiversity computed from audio recordings collected in ecosystems of interest are correlated to the number of bird species recorded during point counts. In particular the ACI shows a significant positive correlation to the number of bird species recorded at point count locations near the recording location. Perhaps surprisingly, we found that the positive correlation between the ACI and the number of birds species identified using the audio recordings was not significant. It is also interesting to note that the location with the lowest ACI, R7, is closest to a major highway which borders the DeKUWC. This location also has the lowest number of species identified using the audio recordings. This points to the effects of major transport highways on biodiversity. A number of authors have investigated the effects of highway noise on bird populations. In [24], Parris *et al.* observed that presence of traffic noise reduced the probability of detecting birds pointing to the possibility of population reduction near roads. In addition, it was observed that certain birds modified their song in presence of traffic noise. In [25], Ware *et al.* used a phantom road experiment to demonstrate the effects of traffic noise on bird populations even without the presence of an actual road. It was found that overall body condition of birds was affected near the phantom road and that birds avoided the area near the phantom road.

In comparing the birds identified during the point counts and those identified using audio recordings, we find that the overlap between the two sets is low. On January 5th 2016, only 12 of the 40 species were identified by both point counts and audio recordings. This is an overlap of 30%. On January 28th 2016, 9 of the 35 species were identified by both point counts and audio recordings. This is an overlap of 25.7%. The low similarity in the results of the point counts and audio recordings could be due to a number of factors including 1) Presence of species which vocalize only rarely 2) Missed identification of bird species vocalizations in the recordings and 3) Possible influence of the vegetation. As has been noted by other authors, point counts and acoustic studies can produce similar or divergent results and this can be influenced by the vegetation in the area of the study [26, 27].

To form an accurate picture of the state of an ecosystem using acoustic recordings, it may be important to build recognizers for particular species known to be common in the ecosystem. This is an approach that has been successfully used in a number of studies such as detecting calls of the Screaming Piha in French Guiana [11] and detecting Nightjar calls in Northumberland, UK [6]. Here we successfully demonstrate the screening of a large audio dataset for calls of the Hartlaub’s Turaco which is a ubiquitous species in montane forests in Kenya with a distinct call. The screening is based on a GMM classifiers trained using MFCCs and simple spectral features. This technique has been successfully used to survey birds that cannot be surveyed using traditional point counts such as nocturnal species for example the European nightjar [6] and Eurasian bittern [7]. In Kenya, the technique applied here can be used to survey the endangered Sokoke Scops Owl without relying on audio playback as is currently done [28].

The monitoring of tropical montane bird species is also important due to the fact that they are particularly vulnerable to the effects of climate change [12]. It is predicted that a number of these birds will lose more than 50% of their range [13]. The work presented here on the automatic detection of the Hartlaub’s Turaco could be used for long-term monitoring of the species. This will ensure that any reduction in the range of the bird is detected early.

In addition to detecting presence of calls in a recording, we are able to detect the number of such calls. This approach can be used to approximate the number of vocalizing individuals in the neighborhood of the recorder. If baseline call rates for a particular species are known, increases in this rate could be indicative of new individuals in the vicinity of the recorder. In addition, the computation of call rates for different species can be used to determine the probability of detection of the species. This approach has the potential to correct for over estimation of the abundance of vocal species.

Acoustic recordings can also be used to estimate the population density of vocalizing species. To do this accurately, distances between the microphones and the vocalizing birds can be estimated using time difference of arrival (TDOA) techniques where the time differences in arrival of an acoustic signal at multiple microphones is used to estimate the distance of the microphones to the sound source [29]. This has been demonstrated in [7] to estimate the spatial distribution of Eurasian bittern.

## Conclusions

This paper has demonstrated the use of acoustic recordings obtained using low cost electronic components in biodiversity monitoring and bird species identification. Using a Raspberry Pi connected to a cheap microphone, audio recordings were obtained from the DeKUWC and used to compute acoustic indices of biodiversity, detect vocalizing bird species and develop an automated bird identification system. We found that acoustic indices of biodiversity computed from these audio recordings were correlated to the number of bird species recorded during point counts. However, we also found that the overlap between the birds identified during the point counts and those identified using audio recordings was low.

This work has demonstrated the use of automatic bird species recognizers to determine the presence of the Hartlaub’s Turaco by recognizing its call. The recognizers used here are based on Gaussian mixture models trained using simple spectral features and MFCCs obtained from the audio recordings. We also demonstrate the use of these GMM classifiers to screen large audio datasets for vocalizations of interest. Building such automatic systems makes screening of these large datasets feasible. For example, there are 331 one minute recordings obtained at recorder location 4, approximately 5.5 hours of audio. We are able to screen each recording in less that 0.5 seconds with the entire dataset screened in approximately three minutes. This represents a considerable time saving.

## Acknowledgments

We would like to thank the Kenya Education Network (KENET) for financial support to purchase recording equipment and all the wardens at the Dedan Kimathi University Wildlife conservancy especially Mr. Rashid Saidi and Mr. Kaindio Kimathi for support during fieldwork.

